# Genome-wide association analysis for resistance to infectious pancreatic necrosis virus identifies candidate genes involved in viral replication and immune response in rainbow trout (*Oncorhynchus mykiss*)

**DOI:** 10.1101/569632

**Authors:** Francisco H. Rodríguez, Raúl Flores-Mara, Grazyella M. Yoshida, Agustín Barría, Ana M. Jedlicki, Jean P. Lhorente, Felipe Reyes-López, José M. Yáñez

## Abstract

Infectious pancreatic necrosis (IPN) is a viral disease with considerable negative impact on the rainbow trout (*Oncorhynchus mykiss*) aquaculture industry. The aim of the present work was to detect genomic regions that explain resistance to infectious pancreatic necrosis virus (IPNV) in rainbow trout. A total of 2,278 fish from 58 full-sib families were challenged with IPNV. Of the challenged fish, 768 individuals were genotyped (488 resistant and 280 susceptible), using a 57K single nucleotide polymorphisms (SNPs) panel Axiom®, Affymetrix®. A genome-wide association study (GWAS) was performed using the phenotypes time to death (TD) and binary survival (BS), along with the genotypes of the challenged fish using a Bayesian model (Bayes C). Heritabilities for resistance to IPNV estimated using pedigree information, were 0.39 and 0.32 for TD and BS, respectively. Heritabilities for resistance to IPNV estimated using genomic information, were 0.50 and 0.54 for TD and BS, respectively. The Bayesian GWAS detected a SNP located on chromosome 5 explaining 18% of the genetic variance for TD. A SNP located on chromosome 23 was detected explaining 9% of the genetic variance for BS. The proximity of Sentrin-specific protease 5 (*SENP5*) to a significant SNP makes it a candidate gene for resistance against IPNV. However, the moderate-low proportion of variance explained by the detected marker leads to the conclusion that the incorporation of all genomic information, through genomic selection, would be the most appropriate approach to accelerate genetic progress for the improvement of resistance against IPNV in rainbow trout.

## INTRODUCTION

Infectious pancreatic necrosis (IPN) is a highly transmissible disease caused by IPN virus (IPNV) which belongs to the Birnaviridae family, genus Aquabirnavirus (RNAds) (Roberts and Pearson, 2005). This virus affects several wild and cultured aquatic organisms. Salmonid species are especially susceptible to IPNV, thus this disease has a great impact on fish farm operations. The mortality levels during an IPN outbreak are influenced by numerous factors; including species, age of the fish, environmental conditions (Dorson and Touchy, 1981), and genetic background, which has been proven to confer resistance to some Atlantic salmon (*Salmo salar*) (Guy *et al*., 2006) and rainbow trout (*Oncorhynchus mykiss*) families (Flores-Mara et al., 2017).

Disease resistance represents a broad term defined as the ability of a host to exert some degree of control over the pathogen life cycle (Bishop and Woolliams, 2014). From a quantitative genetics point of view, disease resistance in fish can be measured using survival data from either field tests or controlled experimental challenges (Ødegård *et al*., 2011, Yáñez *et al*., 2014). In salmonids it has been possible to determine the presence of significant genetic variation for resistance to bacterial (Gjedrem *et al*., 1991; Gjedrem and Gjøen, 1995; Leeds *et al*., 2010; Silverstein *et al*., 2009; Yáñez *et al*., 2013; Yáñez *et al*., 2016; Vallejo *et al*., 2017; Yoshida *et al*., 2018b; Barría *et al*., 2018), parasitic (Glover *et al*., 2005; Kolstad *et al*., 2005; Lhorente *et al*., 2012; Yáñez *et al*., 2013; Ødegård *et al*., 2014), and viral diseases (Guy *et al*., 2006; Henryon *et al*., 2005; Ødegård *et al*., 2007). This implies the possibility of improving, through artificial selection, resistance to various diseases in order to enhance disease control strategies in farmed fish.

The development of genomic technologies has allowed the identification of QTL (quantitative trait loci) or genes involved in the variation of quantitative characteristics (Goddard and Hayes, 2009). The detection of QTL is the starting point for determining functional variants involved in quantitative traits. In addition, this information could be used to accelerate the improvement of traits through the application of marker assisted selection (MAS) or genomic selection (GS) (Yáñez *et al*., 2015). The identification of genes that underlie QTLs can lead to fundamental knowledge of genetic regulation for disease resistance and host-virus interactions in fish (Moen et al., 2009).

QTL for resistance against various diseases have been determined in salmon and trout (Barría et al., 2018; Kjøglum et al., 2006; Ozaki et al., 2007; Correa et al., 2016; Correa et al., 2015; Moen et al., 2015; Houston et al., 2008; Houston et al., 2012). In Atlantic salmon there are some reports of a major QTL for resistance against IPNV in post-smolts, based on data from both field outbreaks (Houston et al., 2008) and experimental challenges (Moen et al., 2009). However, in rainbow trout there is limited information on the molecular genetic bases of resistance to IPNV. Thus, the aim of the present study was to perform a genome-wide association (GWAS) analysis to determine the genetic architecture for IPNV resistance and to identify potential candidate genes involved in the genetic variation of this trait in rainbow trout.

## MATERIALS AND METHODS

### Fish

The population used in this study belonged to the breeding population maintained at Aguas Claras S.A. Experimental tests were conducted at the ATC Patagonia Research Center - Aquainnovo (Puerto Montt, Chile). The fish belonged to 58 full-sib families generated from 58 females and 20 males of rainbow trout from the 2014 year-class. Fertilized eggs from each family were hatched separately. Fish were individually identified using PIT-tags (Passive Integrated Transponders) inserted in the abdominal cavity at an average size of 2g to maintain pedigree traceability. For more details on the description about rearing conditions and population management see Flores-Mara et al. (2017), Neto et al. (2019) and Rodríguez et al. (2018).

### Experimental challenge

Before the experimental challenge, the health condition of 30 random individuals from the full-sib families was evaluated through assessing presence of *Flavobacterium psychrophilum* and IPNV. Fish were challenged with a virulent Sp serotype of IPNV as previously described (Flores-Mara *et al*., 2017). Briefly, fish at an average age of 154 (SD = 15) days, weighing an average of 2.24g (SD = 0.71) were kept in a single 0.25 m^3^ tank under a fresh water recirculation system. Water average oxygen saturation, temperature and salinity during the challenge were 95.74%, 11 °C, and 3.46 ppt, respectively. The virus (CD-AQ03) was isolated using the CHSE-214 cell line at Centrovet Ltda. (Puerto Montt, Chile) from kidney obtained from infected animals from a Chilean Atlantic salmon farm, located in the Xth Region, in November 2014. The isolate was preserved at −80°C until the inoculum preparation using the RTG-2 cell line.

The experimental challenge was performed in two steps; i) intraperitoneal injection of inoculum, at a concentration of 10^7.82^ TCID50/mL, using a quantity of 0.05 mL/fish; and ii) immersion pouring 1.1 L of the inoculum plus 5 L of water into the tank containing 130 L of fresh water, maintained at retained flow at 17 °C for 4 h. After immersion, additional fresh water was introduced at 10 °C, to generate a thermal shock. Mortality was recorded daily and individually recorded. At day 63 the experiment was stopped and all surviving animals were euthanized. The cause of death was confirmed by molecular diagnosis using quantitative real-time reverse transcriptase PCR to determine the presence of IPNV as described by Bowers et al. (2008). Fin clip samples were taken from 768 fish and stored in 98% ethanol at −80°C until genomic DNA extraction. The phenotypic data for resistance to IPNV were obtained from a total of 2,278 fish.

### Genotyping

Genomic DNA was extracted from fin clip samples (n=768 fish) using the commercial kit DNeasy Blood & Tissue Kit (Qiagen) following manufacturer’s instructions. The genomic DNA from each fish was genotyped using a 57K Affimetrix Axiom SNP array developed by Palti et al. (2015). The Affymetrix Analysis Suite AXIOM Software (default parameters) was used to perform the initial quality control (QC) based on clusterization patterns of genotypes. Subsequently, a QC was performed for the genotypes using the R software and the GenAbel package (GenAbel, 2015), using the following parameters: Hardy Weinberg equilibrium (p <1 × 10^−10^), minor allele frequency (> 0.01) and call rate for SNPs and samples (> 0.95). The animals and markers which passed the QC were subsequently used for genomic association analysis.

### Genome-wide association analysis (GWAS)

The resistance phenotypes were defined as the time to death (TD), measured in days with values ranging from 1 to 63, depending on the day of death; and the binary survival (BS), recorded as 1 if the individual died during the challenge and 0 if the individual survived until the end of the trial. To identify the association between SNPs and resistance to IPNV, a linear regression model was used to analyze the TD trait and logistic regression for the binary survival trait (BS). The Bayes C (Habier et al. 2011) analyses were performed using the GS3 software (Legarra et al. 2010). A total of 200,000 iterations were used in the Gibbs sampling, with a burn-in period of 50,000 cycles where results were saved every 50 cycles, totaling 4,000 samples. Convergence and autocorrelation were assessed by visual inspection of trace plots of the posterior variance components. The adjusted model can be described, in matrix notation, as follows: 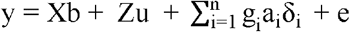, where *y* is the vector of phenotypic records for TD or BS; **X** is an incidence matrix of fixed effects; b is the vector of fixed effect (tagging weight as covariate); **Z** is an incidence matrix of polygenic effects; u is a random vector of polygenic effects of all individuals in the pedigree; g_i_ is the vector of the genotypes for the *i^th^* SNP for each animal; u is the random allele substitution effect of the *i^th^* SNP; a_*i*_, is the random allele substitution effect of the *i_th_* SNP, δ_i_ is an indicator variable (0, 1) sampled from a binomial distribution with such determined parameters that 1% of the markers were included in the model; and *e* is a vector of residual effects. The percentage of the genetic variance explained by each SNP was calculated according to the following formula: 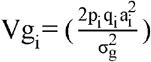 where p_i_ and q_i_ are the allele frequencies for the i-th SNP; ai is the estimated additive effect of the i-th SNP on the phenotype; and is 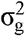 the estimate of the polygenic variance (Lee et al. 2013).

The association of the SNPs with phenotypes were assessed using Bayes factor (BF), which was calculated as follows: 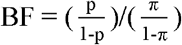, where p is the posterior probability of a SNP to be assigned a non-zero effect and π = 0.99 is the a priori probability of a SNP to be included in the analysis (Kass & Raftery 1995; Varona et al. 2001).

### Candidate genes

The nucleotide sequences surrounding the top ten SNPs that accounted for the largest proportion of genetic variance for each of the IPNV resistance traits were positioned in the most recent version of the rainbow trout reference genome available from NCBI (GenBank assembly Accession GCA_002163495) using BLASTx (Basic Local Alignment Search Tool) (Altschul *et al*., 1997). The presence of annotated genes within 1Mb windows surrounding the top ten SNPs was assessed.

### Data availability

Phenotye data and genotypes are available at the online repository figshare (10.6084/m9.figshare.7725668).

## RESULTS

### Experimental challenge and samples

During the 63 days of the experiment, a maximum number of 23 deaths/day and an average of 6 dead fish per day (SD = 4) were observed. The percentage of accumulated mortality was 13.77%, ranging between 0 to 47.6% for the most and least resistant family, respectively. Out of the 2,278 challenged fish, a selective genotyping strategy was carried out and 768 fish were selected (488 resistant and 280 susceptible). The genotyped fish were selected so that each family was represented, having on average 13.2 (SD = 1.5) fish, with a maximum of 18 and a minimum of 12 fish per family. **Table 1** shows the summary statistics of the animals selected for genotyping.

**Table 1.** General statistics of for the 768 genotyped fish.

| Description | Mean | Standard deviation | Standard error | Minimum | Maximum |
| --- | --- | --- | --- | --- | --- |
| **Weight (g)** | 2.12 | 0.74 | 0.03 | 0.80 | 5.80 |
| **Binary survival** | 0.36 | 0.48 | 0.02 | 0.00 | 1.00 |
| **Time to death** | 51.46 | 13.98 | 0.51 | 13.00 | 62.00 |

### GWAS and candidate genes

A total of 721 individuals and 38,296 SNP passed QC. The average number of SNPs per chromosome was 1,277 (SD = 384) and varied between 594 and 1,818. The variance components and heritabilities estimated using genomic information are shown in Table 2. Heritabilities were high and similar between the two trait definitions (0.50 0.06 and 0.54 ± 0.09 for TD and BS, respectively). The top SNP for both traits explained 18.89% and 9.31% of the genetic variance for TD and BS, respectively. According to Vidal et al. (2005), a BF from 3 to 20 is suggestive of linkage and BF from 20 to 150 indicates linkage with the SNPs and the trait under investigation. In this study, we focused the results and discussion on the SNPs with a BF greater than 20. For TD, the top four SNPs that presented a BF greater than 20 are located on chromosomes 5 (AX-89921775), 13 (AX-89964133), 21 (AX-89928391) and 30 (AX-89972475). The sum of these SNPs explained 32.99% of all genetic variance. For the BS trait only the top one SNP presented a BF greater than 20, and it is located on chromosome 23 (AX-89938762) and explained 9.31% of the genetic variance for this trait. The most relevant genes next to the top ten SNPs that explained the highest proportion of genetic variance for each trait are shown in **Table 3**.

**Table 2.**
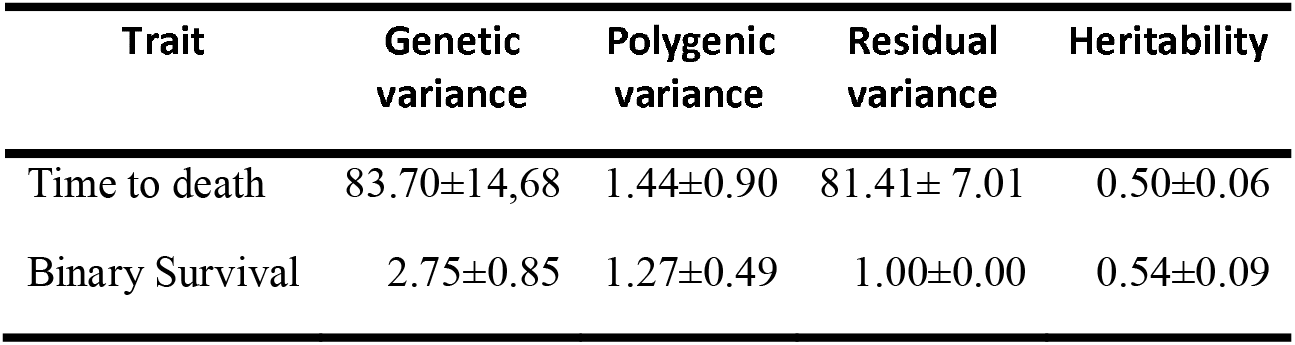
Mean ± standard deviation of variance components explained by whole genome SNP markers for time to death and binary survival for rainbow trout using Bayes C.

| Trait | Genetic variance | Polygenic variance | Residual variance | Heritability |
| --- | --- | --- | --- | --- |
| Time to death | 83.70±14,68 | 1.44±0.90 | 81.41± 7.01 | 0.50±0.06 |
| Binary Survival | 2.75±0.85 | 1.27±0.49 | 1.00±0.00 | 0.54±0.09 |

**Table 3.**
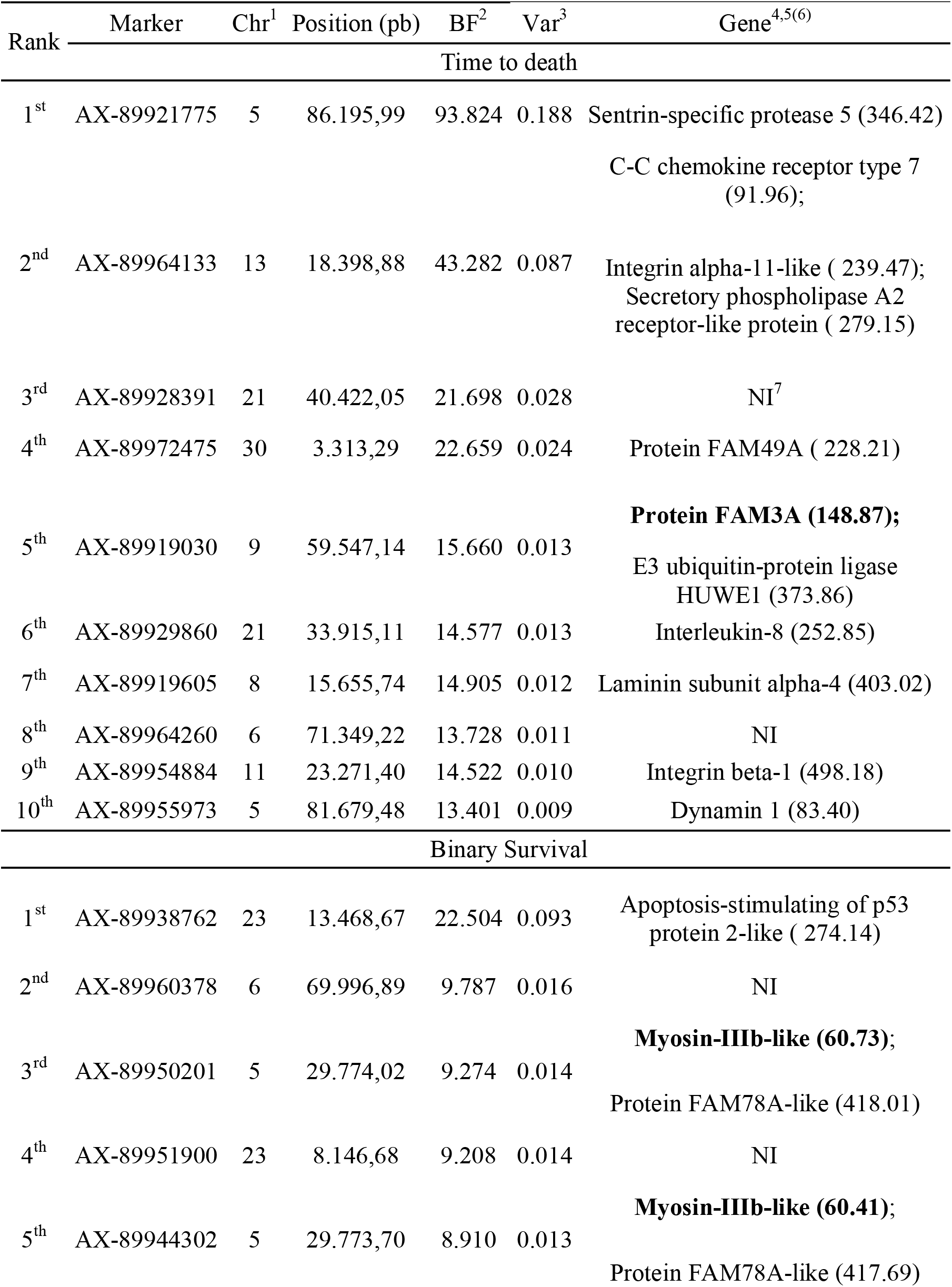
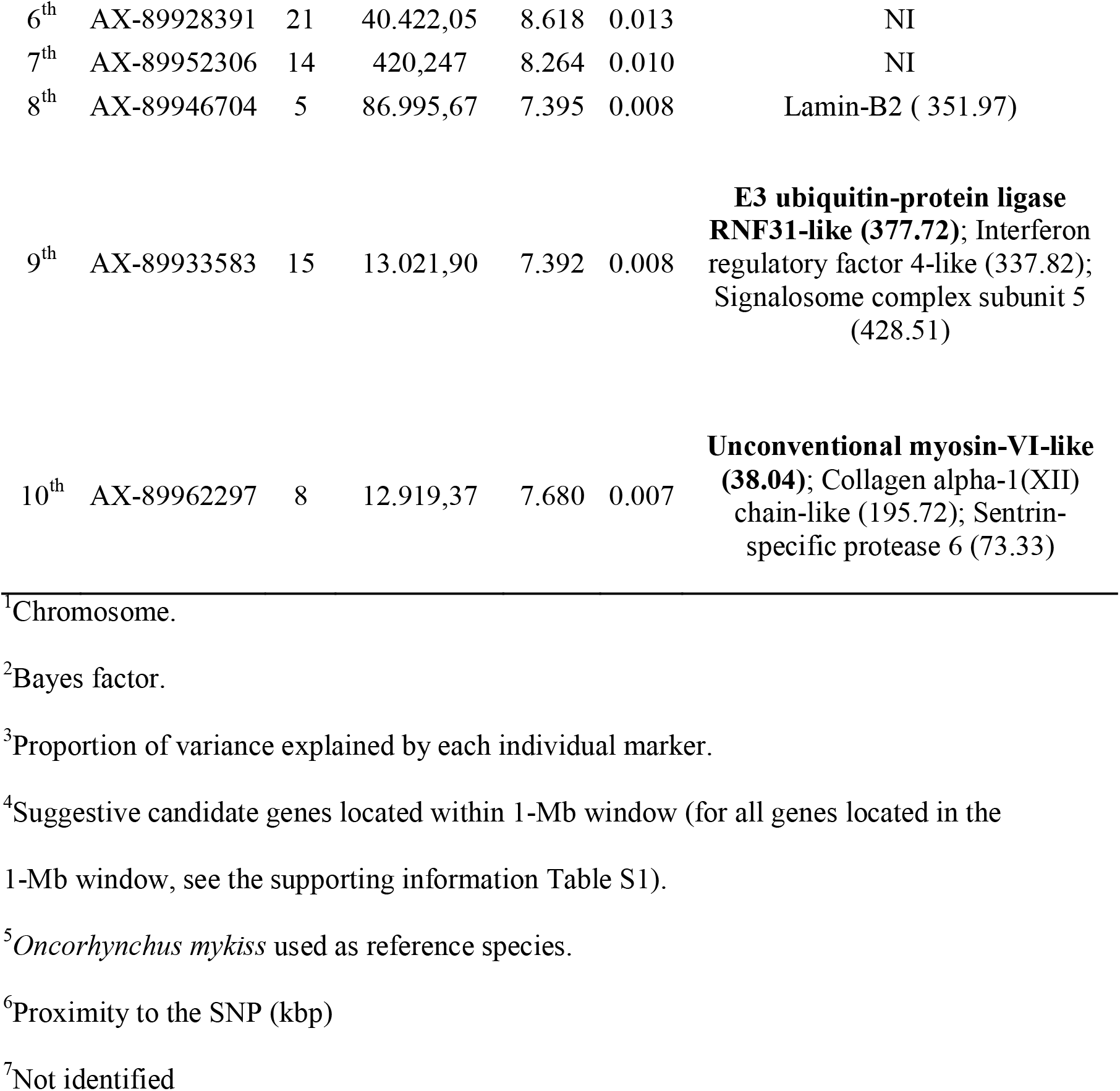
Top 10 SNPs associated with resistance to IPNV in rainbow trout using Bayes C model.

## DISCUSSION

The mortality rate in IPNV challenge experiments is highly variable and depends on several factors the virus strain, the population evaluated, the size of the fish, environmental conditions, and others. For example, mortality rates over 40% have been reported for Atlantic salmon (Robert and Pearson, 2005; Houston *et al*., 2008). Other authors have reported mortality rates of 70.5% and 77.8% for the same disease and species (Moen *et al*., 2009), while in rainbow trout mortality rates range from 36% to 54% (Ozaki *et al*., 2007). The level of mortality makes it important to study traits in rainbow trout that can be selected for, to create IPNV-resistant fish.

Some studies have shown significant pedigree-based heritabilities for IPNV resistance in rainbow trout and Atlantic salmon (Flores-Mara *et al*., 2017; Guy *et al*., 2006; Storset *et al*., 2007) using survival time as the trait definition. For instance, in Atlantic salmon, heritability values ranging between 0.26 and 0.55 have been reported for resistance to IPNV, measured as binary survival and analyzed by threshold models (Wetten *et al*., 2007; Kjøglum *et al*., 2008; Guy *et al*., 2009; Gheyas *et al*., 2010; Houston *et al*., 2010). Previous studies have reported pedigree-based heritability values of 0.39 and 0.4 for time to death and 0.35 for binary survival in the same population of IPNV-infected rainbow trout (Flores-Mara *et al*., 2017;Yoshida et al. 2018a). Moreover, heritability values of 0.24 and 0.25 for day to death and binary survival, respectively, have been reported (Yoshida *et al*., 2018a) using a combination of both the pedigree relationship matrix (A) and the genomic matrix (G), also called the H matrix (Aguilar et al. 2010). The heritability values calculated in the present study using genomic information are higher than values previously reported in rainbow trout and indicate that selection to improve resistance to IPNV is feasible. The proportion of genetic variance explained by the SNP with the largest effect was larger for trait TD than the SNP identified for trait BS. The BF indicated that four SNP were associated with TD and one for BS.

Previous studies in Atlantic salmon have determined two significant QTLs for resistance against IPNV. The most significant, explaining 29% and 83% of the phenotypic and genetic variance, respectively, was identified on chromosome 26 (Houston *et al*, 2008; 2012). This QTL was confirmed in an independent Atlantic salmon population from Norway (Moen *et al*., 2009). A recent study showed that resistance to IPNV in Atlantic salmon is determined by a locus of major effect, which is probably influenced by variations in a gene coding sequence for the epithelial cadherin gene (Moen *et al*., 2015). However, in the present study focusing on rainbow trout, no QTL were found to co-localize with the regions previously associated with resistance to IPNV in Atlantic salmon (Phillips *et al*., 2009). Concordantly, the QTLs reported by Ozaki *et al*. (2001; 2007) in rainbow trout are not found in homologous regions in Atlantic salmon (Danzmann *et al*., 2005; Phillips *et al*., 2009). In rainbow trout, Ozaki *et al*. (2001) found 2 QTLs associated with resistance to IPNV using the linkage map elaborated by Sakamoto *et al*. (2000) and based on microsatellite markers. The first QTL was found in linkage group 3 while the second QTL was identified in linkage group 22, each of which explains about 17% of the phenotypic variance (Ozaki *et al*., 2001). The same QTLs were confirmed in a subsequent study (Ozaki *et al*., 2007), in which another significant QTL was detected in linkage group 12. Based on the linkage map developed by Palti *et al*. (2012) these significant markers were located on chromosome 14 and 16, respectively (Zhi-Liang *et al*., 2016).

The SNP that explained the greatest proportion of genetic variance for TD in the present study (AX-89921775) is located on chromosome 5 in a region that contains an important gene that encodes *Sentrin-specific protease 5*. In mammals this protein participates in the SUMO pathway (*small ubiquitin-like modifier*) and its function is mainly related to the activity of isopeptidase (elimination of SUMO chains) (Pinto *et al*., 2012). SUMO is involved in the regulation of biological processes that are key to viral replication, including genetic transcription, cell cycle, apoptosis, intracellular and intranuclear trafficking, and protein stability.

Resistance to infection can be the result of a more robust host immune response to the invading agent. In this study we identified SNP located near several genes that encode proteins whose function is to control inflammation as well as other immune-related responses. Close to SNP AX-89964133, located on chromosome 13, there is a gene that encodes *Secretory phospholipase A2 receptor-like protein*. Phospholipases A2 are enzymes released in plasma and extracellular fluids during inflammatory diseases. This protein induces the release of the pro-inflammatory cytokines TNF-α and IL-6 depending on the concentration of protein (Granata *et al*., 2005). These cytokines participate in the differentiation of B lymphocytes and activation of cytotoxic T lymphocytes (Reyes-Cerpa et al., 2012). In addition, TNF-α participates in the recruitment of immunoglobulins important for controlling infection.

One of the key host responses against viral agents is mediated by interferon (IFN) proteins which activate the cell antiviral state, thus IFN plays a major role in the activation of defense mechanisms against viral infection in vertebrates (Robertsen, 2006). After the specific recognition of viral antigens, the IFN response is induced through the activation of interferon regulatory factor (IRF). Importantly, we identified a SNP associated with BS, which is located in a region that encodes *interferon regulatory factor 4-like* (*irf4*) (SNP AX-89933583; chromosome 15). This protein is important in both the regulation of interferons in response to virus infection and the regulation of inducible genes by interferon. In fact, in mammals *irf4* plays a central role as a TH1 regulator (Mahnke et al., 2016), a fundamental cell-mediated process for the host antiviral response. Furthermore, in Atlantic salmon IPNV-susceptible families have reported a higher expression of *ifnα* and *ifnγ* compared to the resistant phenotype (Reyes-López et al., 2015). Therefore, the SNP located in the region encoding for *IRF-4-like* might provide additional evidence for the role of interferon in salmonid IPNV-resistance. Other studies have revealed that IRF also plays a critical role in other biological processes, such as metabolism (Zhao *et al*., 2015). This suggests that *irf4* could play a role in the IPNV-resistance phenotype through other pathways not directly associated with the immune response.

SNP AX-89933583 is also located in a gene region that encodes for *signalosome complex subunit 5-like*. This protein is involved in the regulation of nuclear factor kappa B (NF-kB) one of the most relevant transcription factors involved in the control of several pro-inflammatory proteins (Adcock and Caramori, 2001). This SNP is also located in the gene region that encodes *E3 ubiquitin-protein ligase RNF31-like*, which in mammals, plays a key role in NF-kB activation (Tokunaga et al., 2009; Kirisako et al., 2006). Together, these SNPs could affect the gene function of important mediators of gene expression and activation of the pro-inflammatory response.

The activation of the immune response also requires the expression of genes responsible for the activation and recruitment of immune cell populations to the site of infection. In our study, SNP AX-89929860 was located in a gene region that codes for *Interleukin-8*, also named *IL-8. Interleukin-8* is a member of the CXC chemokine subfamily directly associated with the pro-inflammatory response because it attracts different immune cell populations including neutrophils, T lymphocytes and basophils (Reyes-Cerpa et al., 2012). Importantly, a gene expression increase for *Interleukin-8* was observed in symptomatic IPNV-infected head kidney trout but not in fish with persistent asymptomatic infection (Reyes-Cerpa et al., 2014), suggesting that SNP AX-89929860 is associated with resistance to IPNV because of its potential effect on the inflammatory response.

Once the host immune response is activated, one of the crucial processes is leukocyte redistribution to the site of infection, which involves cell binding to endothelia and trans endothelial migration. In our study, several genes related to this process were located in regions where SNPs associated with IPNV resistance were identified. Among them, *C-C chemokine receptor type 7* (which is close to SNP AX-89964133) is described as a potent leukocyte chemotactic receptor, also responsible for directing the migration of dendritic cells (DCs) to the lymph nodes, where these cells play an important role in the initiation of the immune response. Recently, it has been described that *C-C chemokine receptor type 7* controls cytoarchitecture, the rate of endocytosis, survival, migration speed and DC maturation (Sánchez-Sánchez *et al*., 2006). There are no studies to date that evaluated the effect of *C-C chemokine receptor type 7* in salmonids infected with IPNV.

Located close to the SNP AX-89919605 is *Laminin subunit alpha 4* which is also associated with cell adhesion, differentiation, and cell migration. SNP AX-89954884 is located near *integrin beta 1;* which is a membrane receptor involved in cell adhesion and recognition in a variety of processes including the immune response. Importantly, the *integrins* has been described as a receptor for the pro-inflammatory cytokine IL-1β, whose binding is essential for the IL-1β signaling (Takada et al., 2017); suggesting that this SNP could influence the pro-inflammatory IPNV-infected fish response.

Previous reports in Atlantic salmon have shown clear differences in the gene expression pattern between IPNV-susceptible and –resistant phenotypes showing differences in ubiquitin-dependent and apoptosis processes (Reyes-López et al., 2015; Robledo et al., 2016). In our study two SNPs (AX-89919030; AX-89938762) were located in regions that contain genes that encode for *E3 ubiquitin-protein ligase HUWE1* and *Apoptosis-stimulating of p53 protein 2-like*, respectively. *E3 ubiquitin-protein ligase HUWE1* mediates the proteasomal degradation of target proteins, providing more evidence that ubiquitin-dependent processes could be important factors in IPNV disease resistance. The role of *HUWE1* in proteasomal degradation of target proteins opens the possibility that other processes associated with this function, like vesicular trafficking, could also be involved with susceptibility to IPNV. In support of this hypothesis, we found SNP AX-89955973 close to the gene region for *dynamin 1*, a member of the GTP-binding proteins involved in clathrin-mediated endocytosis and other vesicular trafficking processes. Further studies to evaluate the functional role of these processes in the salmonid IPNV resistance phenotype remains to be elucidated.

In IPNV-susceptible Atlantic salmon, it has been shown that the modulation of myosin-related processes are linked to the abnormal swimming behavior observed in IPNV-affected fish (Robledo et al., 2016). Interestingly, two SNP on chromosome 5 (AX-89950201 and AX-89944302) and one on chromosome 8 (AX-89962297) were located near *myosin-IIIb-like* (Chromosome 5) and the *unconventional myosin-VI-like* (Chromosome 8) genes.

The results obtained from this study indicate that the QTLs involved in IPNV resistance contribute to a moderate-low proportion of the variance of this trait, so the implementation of this information in MAS is probably not the most efficient approach (Correa *et al*., 2015). Our data indicates that molecular information that results from genotyping high-density panels of SNPs can be efficiently incorporated into breeding programs to accelerate the development of IPNV-resistant strains through the application of genomic selection (Goddard and Hayes, 2007).

## CONCLUSIONS

In conclusion, this is the first work reporting the detection and position of QTL involved in IPNV resistance in rainbow trout using SNP type markers. Resistance to IPNV can be described as a moderately oligogenic trait since there are probably several loci involved; each with a moderate-low effect. Potential candidate genes have been identified close to the associated SNPs, whose biological role in the fish immune response suggest they could be involved in the mechanisms of resistance against IPNV.

**Figure 1.**
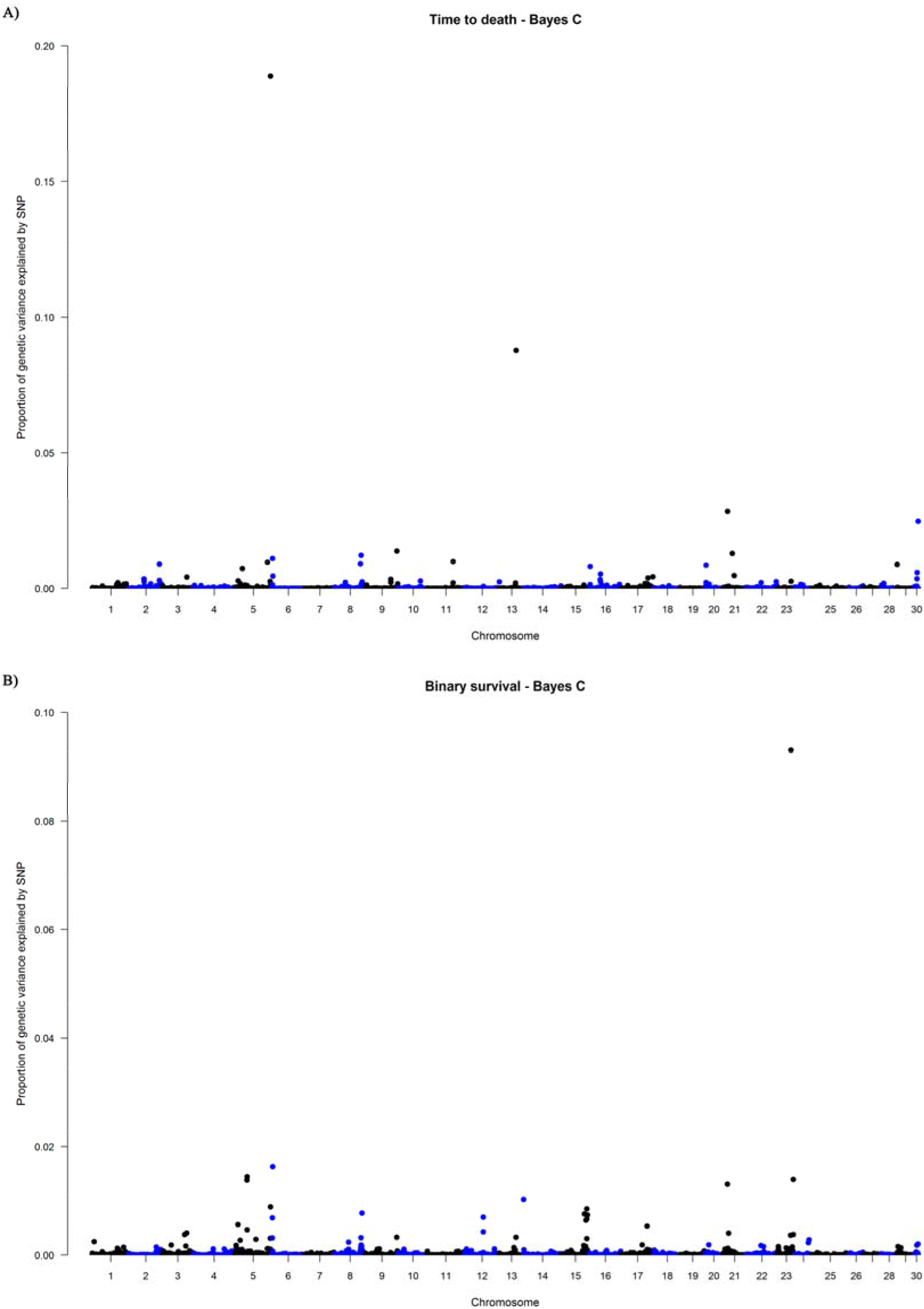
Manhattan plot of variance explained by each SNP using Bayes C model. (A) Time to death. (B) Binary survival.

## Acknowledgments

The authors would like to thank the company Aguas Claras S.A. for providing the fish and data used in the present study. This study has been partially funded by a CORFO grant (12PIE-17669), Government of Chile. Francisco H. Rodríguez and Raúl Flores-Mara would like to thank the grant Presidente de la República of the Government of Peru.

## Authors’ contributions

FHR performed the analysis and wrote the inicial version of manuscript. RFM contributed to the analysis and contributed to discussion and writing. GMY performed the GWAs analysis and contributed to discussion and writing. AB contributed to the analysis, discussion and writing. AMJ performed the DNA extract. JPL contributed with study design. FRJ contributed with results interpretation, discussion and writing. JMY conceived and designed the study; contributed to the analysis, discussion, writing and supervised work of FHR. All authors have reviewed and approved the manuscript.

## Animals ethics approval

Challenge and sampling procedures were approved by the Comité de Bioética Animal from the Facultad de Ciencias Veterinarias y Pecuarias, Universidad de Chile (Certificate N° 17,019–VET-UCH).

